# *DemoTape*: Computational demultiplexing of targeted single-cell sequencing data

**DOI:** 10.1101/2024.12.06.627152

**Authors:** Nico Borgsmüller, Jack Kuipers, Johannes Gawron, Marco Roncador, Marcel Pohly, Erkin Acar, Thi Huong Lan Do, Stefanie Reisenauer, Mirjam Judith Feldkamp, Christian Beisel, Thorsten Zenz, Andreas Moor, Niko Beerenwinkel

## Abstract

**Background:** Single-cell sequencing can provide novel insights into the understanding and treatment of diseases. In cancer, for example, intratumor heterogeneity is a major cause of treatment resistance and relapse. Although technological progress has substantially increased the throughput of sequenced cells, single-cell sequencing remains cost and labor-intensive. Multiplexing, i.e., the pooling and subsequent joint preparation and sequencing of samples, followed by a demultiplexing step, is a common practice to reduce expenses and confounding batch effects, especially in single-cell RNA sequencing.

**Results:** Here, we introduce *demoTape*, a computational demultiplexing method for targeted single-cell DNA sequencing (scDNA-seq) data based on a distance metric between individual cells at single-nucleotide polymorphisms loci. To validate *demoTape*, we sequence three B-cell lymphoma patients separately and multiplexed on the Tapestri platform. We find similar genotypes, clones, and evolutionary histories in all three samples when comparing the individual with the demultiplexed samples. Using the three individually sequenced samples, we simulate multiplexed ground truth data and show that *demoTape* outperforms state-of-the-art demultiplexing methods designed for RNA sequencing data. Additionally, we demonstrate through downsampling that the inferred clonal composition remained largely stable for samples with fewer cells despite the inevitable loss in resolution of low-frequency clones.

**Conclusions:** Multiplexing and subsequent genotype-based demultiplexing of scDNA-seq will reduce costs and workload, eventually allowing the sequencing of more samples. This will open new possibilities and accelerate the investigation of biological questions where cellular heterogeneity on the genomic level plays a crucial role.

## 1 Introduction

With the advent of high-throughput sequencing technologies, DNA and RNA data increased steadily in quality and quantity. The rapid decrease in cost and labor per sequenced sample has enabled the deciphering of thousands of human genomes [1, 2] and hundreds of mammalian species [3]. With the development of droplet-based microfluidic systems, it is now possible to sequence thousands of single cells at reasonable cost and in reasonable time [4, 5] and, additionally study multiomics questions at single-cell resolution.

The increase in data led to a better understanding of diseases with genomic origin. Especially in cancer, intra-tumor heterogeneity (ITH) was identified as a key factor driving clonal evolution and causing treatment resistance [6, 7]. In B-cells, for example, single-cell data revealed that therapeutic pressure increased the complexity of the clonal composition, resulting in treatment resistance [8]. Accordingly, various methods were proposed to infer the clonal composition within a tumor, initially from bulk [9, 10] and, with increasing resolution, from single-cell DNA sequencing (scDNA-seq) data [11, 12, 13].

Although the quantity of single-cell data increased overall, the focus has mainly been on RNA data [14]. scDNA-seq is more cost- and labor-intensive as it relies on amplifying the DNA molecules per cell above the detection threshold, making especially whole-exome and whole-genome sequencing expensive [15]. Novel, target-based approaches like the Tapestri platform [16] amplify only a panel of predefined sites and sequence these in thousands of cells with high accuracy [17].

In general, the information content of a sequencing experiment grows with the number of cells analyzed, but it eventually saturates. Consequently, it is common practice in scRNA-seq to pool (multiplex) multiple samples for joint preparation and sequencing, leading to fewer cells per sample but also decreased expenses and batch effects [18]. For subsequent demultiplexing, i.e., separating cells into their original samples, approaches relying on cell hashing via biochemical assays are often utilized [19, 20] but also computational tools using single-nucleotide polymorphisms (SNPs) as sample markers can be used [21, 22]. However, genotype-based demultiplexing of scRNA-seq data is challenging due to the sparse SNP data, as SNP inference relies strongly on sufficient expression levels [23]. scDNAseq data, in contrast, allows the reliable calling of SNPs which should be sufficient to tell multiplexed patients with distinct SNP profiles apart. Doing so without additional preparation steps and chemicals would reduce the cost and time required per sample. However, droplet-based approaches like the Tapestri platform tend to produce cell doublets (two cells sequenced together) which might distort the called SNPs and hamper the correct assignment of cells to patients [24].

Here, we introduce *demoTape*, a computational demultiplexing method for targeted scDNA-seq data based on the genotype distance between individual cells. Specifically, *demoTape* uses a novel distance metric based on the read counts at SNP loci, leveraging the deep read coverage at the targeted sites. To validate *demoTape*, we sequenced three B-cell lymphoma patients (one mantel cell lymphoma and two marginal zone lymphomas) separately and multiplexed on the Tapestri platform with a custom DNA target panel. Using these data, we demonstrate that *demoTape* outperforms state-of-the-art scRNA-seq methods in accuracy and that the inferred clonal composition remains largely stable when decreasing the number of cells. We then applied *demoTape* to the multiplexed sample, assigning cells back to patients and confirming the assignment through clonal, patient-specific SNPs.

## 2 Results

### 2.1 Overview of *demoTape* and the sequencing data of three lymphoma patients

To computationally demultiplex pooled samples from a joint sequencing run, *demoTape* requires a matrix containing the read counts at all called SNP loci and the number of multiplexed samples as input. First, *demoTape* calculates the pairwise distances between all cells based on the observed reads, creates a cell dendrogram through hierarchical agglomerative clustering, and uses this dendrogram to determine initial clusters. Next, it identifies clusters corresponding to doublets from different samples, checks these for plausibility, and increases the cluster number if not enough plausible doublet clusters are identified. Finally, it merges added clusters (if any) until the cluster number equals the number of pooled samples (Fig. 1a). Section 5.1 describes *demoTape* in greater detail.

**Figure 1:**
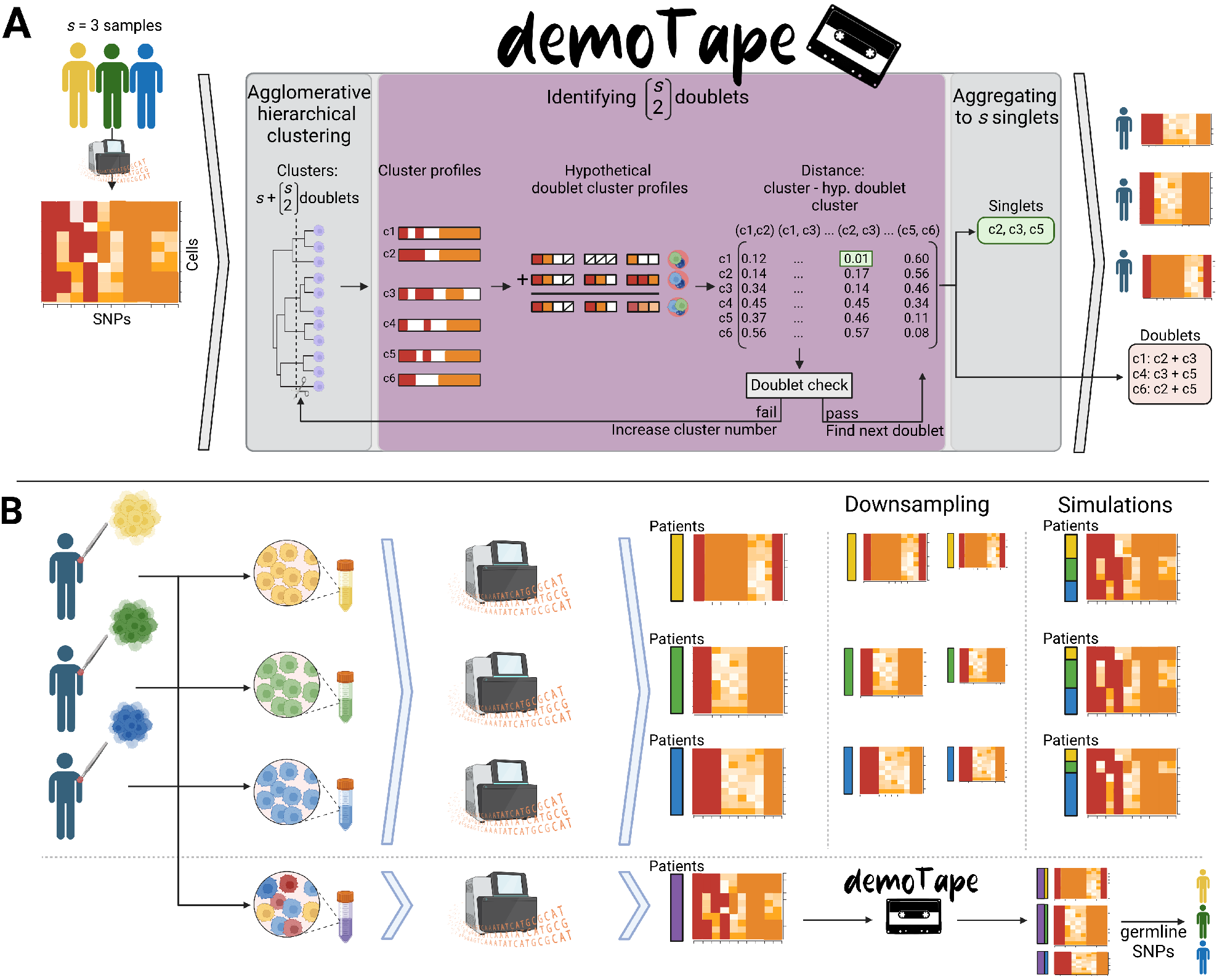
Overview of *demoTape* and conducted experiments. **A)** Given a read count matrix and the number of pooled samples, *demoTape* calculates pairwise distances between cells based on the observed reads, applies hierarchical agglomerative clustering to obtain cell clusters, and calculates corresponding read count profiles per cluster. Next, *demoTape* computes profiles for hypothetical doublets and uses these to identify potential doublet clusters. If not enough doublet clusters are identified, the cluster number is increased. Subsequently, the remaining clusters are aggregated to match the number of pooled samples. **B)** We sequenced the DNA of three patients individually and multiplexed on the Tapestri platform. Next, we synthetically multiplexed cells from the patient samples to benchmark the accuracy of *demoTape* and downsampled the number of cells in the patient samples to evaluate the effect of having fewer available cells per sample on the downstream analysis. Finally, we applied *demoTape* to the multiplexed sample, compared the results to the individual patient samples, and assigned demultiplexed clusters back to patients.

To assess the performance of *demoTape*, We sequenced three lymphoma patient samples individually (S1, S2, and S3) and multiplexed (MS1) on the Tapestri platform (Section 5.4). After processing and applying default filters (Section 5.5), S1 contained 3732 cells and 52 SNPs, S2 contained 7911 cells and 37 SNPs, S3 contained 3677 cells and 40 SNPs, and MS1 contained 4314 cells and 84 SNPs (Fig. 4a). In total, 20 SNPs were called in all four samples, and 2, 1, and 3 SNPs were called only in S1, S2, and S3, respectively. MS1 contained 15 SNPs not called in the individual samples, out of which 13 showed a low variant allele frequency (VAF), i.e., the average VAF at that locus, weighted by the read depth per cell, was less than 20 %. The average VAF of the remaining two SNPs detected only in MS1 were 42 % and 50 %.

Using the separately sequenced samples, we generated computationally multiplexed samples with known ground truth, allowing direct evaluation of *demoTape*. Additionally, we downsampled the cells of the individual samples to assess the effect on the inferred clonal composition. Finally, we analyzed and compared the individual samples with the multiplexed one and assigned the demultiplexed clusters to patients (Fig. 1b).

### 2.2 Distance-based multiplexing is reliable on simulated data

To simulate multiplexed samples with known ground truth, we sampled single cells from S1, S2, and S3, generated doublets consisting of cells from different samples, and filtered loci and cells of the synthetic multiplexed sample (see Section 5.2). In total, we simulated 48 datasets with different mixing ratios and doublet rates and repeated each simulation 30 times. As even mixing ratios cannot be assumed, along with equal ratios of 33:33:33% we simulated all possible combinations of the uneven ratios of 20:40:40% (three combinations), 20:30:50% (six combinations), and 10:30:60% (six combinations). For all mixing ratios, we also simulated doublets with rates of 10, 20, and 30 %. As no methods tailored explicitly at scDNA-seq exist, to the best of our knowledge, we ran souporcell [22], scSplit [21], and Vireo [25] for comparison, three state-of-the-art methods for demultiplexing scRNA-seq data based on genotypes. We evaluated the accuracy of cell assignments, i.e., the fraction of cells assigned to the correct sample or, for doublets, to a general doublet cluster, using the V-measure [26]. We used a general doublet cluster instead of the three specific ones (S1+S2, S1+S3, and S1+S2) for a fair comparison, as scSplit only infers doublets but not their composition.

In all simulated cases, except the ones with even mixing ratios, Vireo performed worst, mainly resulting from a large fraction of cells that Vireo could not assign to any sample or doublet. With even mixing rations, scSplit performed worst and second worst in all other cases. *DemoTape* and souporcell performed equally well with V-measures close to 1, indicating nearly perfect demultiplexing, for even mixing ratios or slightly uneven mixing ratios of 20:40:40%, independent of the simulated doublets. In all cases, *demoTape* performed slightly better; the median V-measure difference between the two methods was smaller or equal to 0.01 (Fig. 2a and b). In simulation scenarios with 20:30:50% and 10:30:60% mixing ratios, *demoTape* outperformed souporcell (Fig. 2c and d). Especially at high doublet rates, souporcells V-measure decreased to 0.9 (10:30:60% mixing ratio with 30 % doublets), whereas *demoTape’s* demultiplexing remained accurate with a median V-measure of over 0.995.

**Figure 2:**
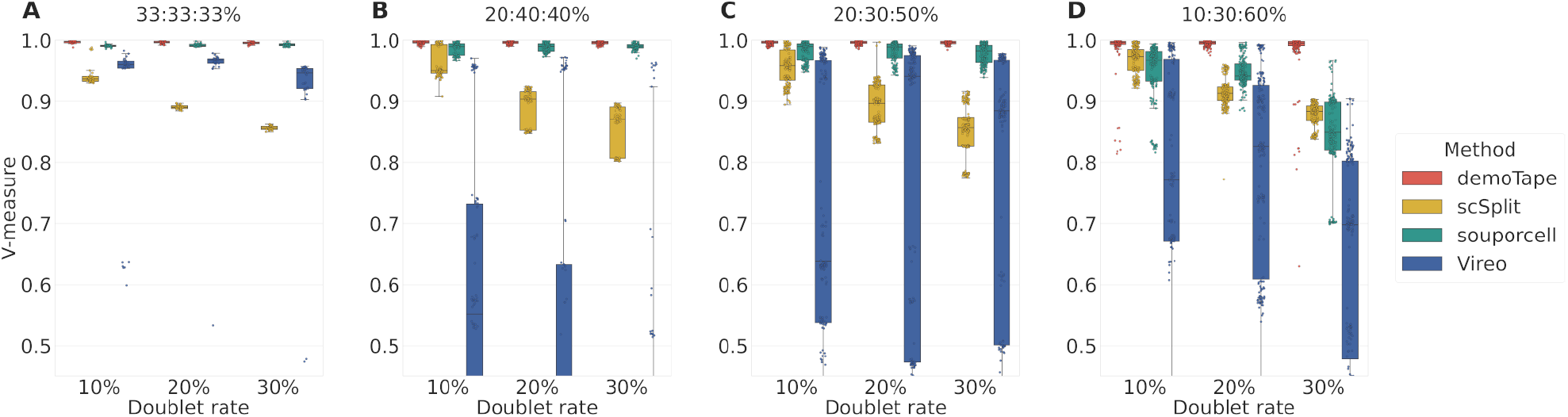
V-measure of the genotype-based demultiplexing methods *demoTape*, scSplit, souporcell, and Vireo on simulated data from three samples, for different mixing ratios and doublet rates of 10, 20, and 30%. Each box summarizes 30 simulations. The mixing ratios varied from uniform 33:33:33% (**A**), to slightly uneven 20:40:40% (**B**), to uneven 20:30:50% (**C**), to highly uneven 10:30:60% (**D**).

Our simulations showed that *demoTape* outperformed the scRNA-seq methods, achieving the highest demultiplexing accuracy on all simulated datasets. For all simulated datasets, its median V-measure was above 0.981, and its average V-measure was 0.958.

### 2.3 Clonal information remains consistent with smaller cell numbers

To assess the effect of the cell number on the inferred clonal composition, we downsampled the number of cells in S1, S2, and S3 without replacement from the total number of cells to 2000, 1000, and 500 cells; for S2 (7911 cells), we additionally downsampled the cell number to 4000 cells. We repeated each downsampling ten times. Then, we processed the downsampled datasets, filtered uninformative, i.e., germline, SNPs (Section 5), and ran the three state-of-the-art scDNA-seq clustering methods SCG [11], BnpC [12], and COMPASS [13]. SCG and BnpC infer clusters with their corresponding genotype profile and outperformed other scDNA-seq clustering methods in a recently published benchmarking study [27]. COMPASS has been designed especially for Tapestri data and infers a mutation tree, where nodes represent clusters with a certain genotype profile, and cells are assigned to these nodes. We ran SCG and COMPASS with doublet rates of 8 and 25 %, the former being the value reported by MissionBio, and the latter being close to the observed rate in MS1. BnpC does not model doublets explicitly. To measure the effect of the downsampling on the inferred clusters, we applied the three clustering methods ten times to each complete sample, containing all cells, and took the inferred clusters as ground truth. Then, we calculated the average adjusted Rand index (ARI) [28] between the clusters inferred from the downsampled and the complete datasets, for each clustering method, where higher values indicate a higher similarity.

For S1, the number of reported SNPs decreased steadily from 18 in the complete sample to an average of 6 in samples with 500 cells, with the highest decrease from 13 to seven appearing from 1000 to 500 cells (Fig. 3A). The median number of inferred clusters was relatively stable at three (± 0.5) for BnpC and between one and two for SCG, independent of the doubled rate (Fig. 3D). Remarkably, the median number of clusters inferred by SCG on the full dataset was 1 and 1.5 (8 and 25 % doublets, respectively), indicating that SCG was unable to identify meaningful clusters. On the complete sample, SCG’s median ARI was 0.8/0.4 (8*/*25 % doublets), and BnpC’s was 0.66, revealing a low clustering consistency when run multiple times on the same data (Fig. 3G). The unchanged ARI of 0.8 for SCG runs with 8 % doublets was due to the inference of only one cluster in most runs (especially in 8/10 ground truth runs), except for 500 cells, where 2 clusters were inferred (ARI = 0.2). For SCG runs with 25 % doublets, 1.5 clusters were inferred on the complete sample, but only one cluster in most subsample runs, leading to an ARI of 0.5. BnpCs ARI decreased steadily with smaller sample sizes to 0.18 (500 cells). COMPASS, in contrast, always inferred the same six clusters on the complete sample (ARI = 1), a median of six clusters on samples with 2000 cells (ARI = 0.86/0.92, for 8*/*25 % doublets), and five and three clusters for 1000 and 500 cells, respectively. Accordingly, its ARI also declined, with the highest decrease occurring at 500 cells (0.55/0.66 to 0, for 8*/*25 % doublets).

**Figure 3:**
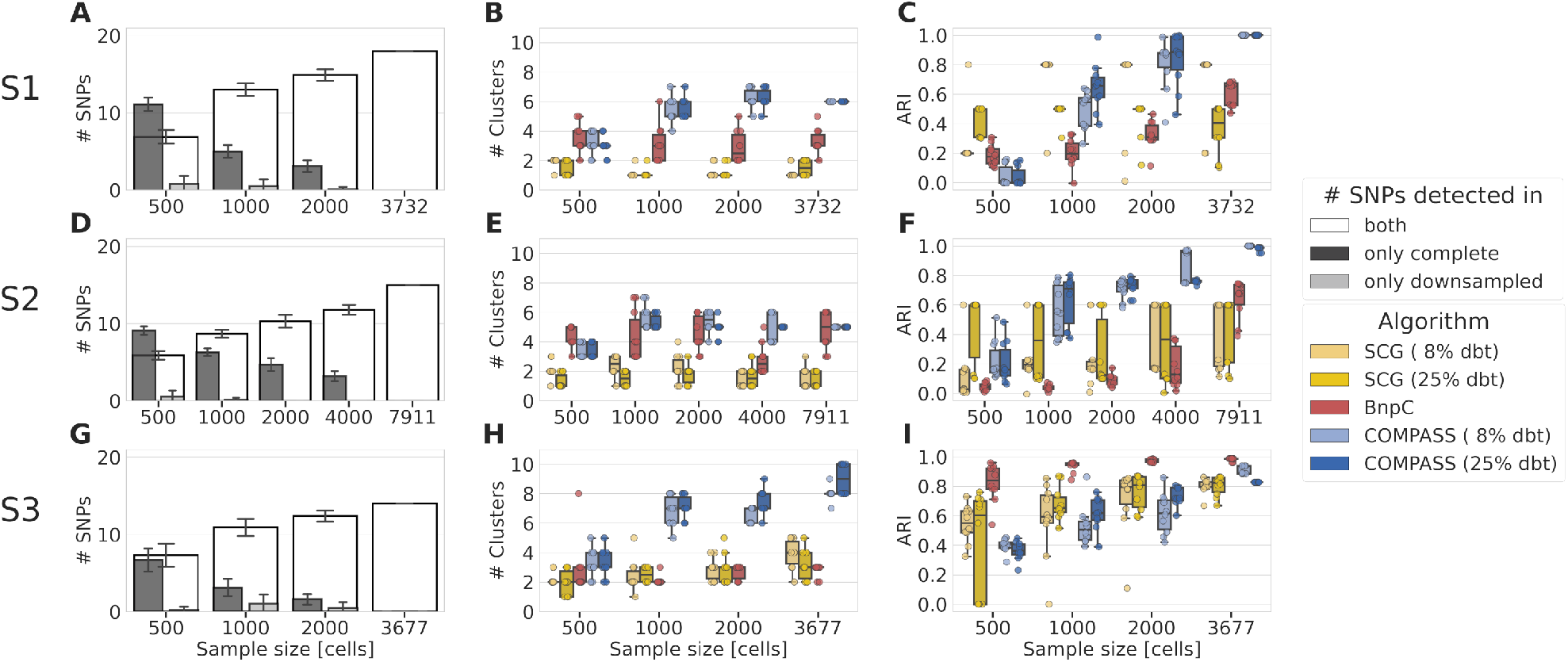
Results of the downsampling simulations (*n* = 10 per sample size) for patients S1 (first row), S2 (second row), and S3 (third row). The first column (**A, D**, and **G**) shows the number of reported SNPs and the overlap of smaller sample size runs with the complete sample. The second column (**B, E**, and **H**) indicates the number of inferred clusters, and the third column (**C, F**, and **I**) the Adjusted Rand Index (ARI) of the clustering algorithms BnpC, SCG, and COMPASS. As ground truth for the ARI calculation, we used the clusters inferred on the complete data containing all cells (last value on the x-axis). Higher ARI values indicate a higher clustering consistency.

In patient S2, fewer informative SNPs were reported, although more cells were sequenced. Of the 15 SNPs reported in the complete sample, 12 were found in the samples with 4000 cells and six in the samples with 500 cells (Fig. 3B). Here, equally high decreases (three reported SNPs fewer) occurred when changing the sample size from the complete sample to 4000 cells and from 1000 to 500 cells. Again, the median number of clusters inferred by SCG on the complete sample was one and remained between one and three on smaller sample sizes (Fig. 3E), resulting in ARI values below 0.6 in all runs (Fig. 3F). BnpC inferred five clusters (median) on the complete sample and four on samples with fewer cells, except for 4000 cells, where the median cluster number was 2.5. Its ARI declined from an initial 0.68 to 0.13 for 4000 cells and remained low (<0.1) for smaller sample sizes. COMPASS inferred five clusters on the complete sample and reported, when run with a doublet rate of 25 %, the same number for all sample sizes, except for 500 cells, where it dropped to three. Running COMPASS with 8 % doublets resulted in higher variations in the cluster number, ranging from four to 5.5 (median) for all sample sizes greater than 500 cells and three for samples with 500 cells. On the complete sample, its ARI was 1.0/0.99 (doublet rate) again, decreasing steadily with a smaller sample size. For 8 % doublets, it remained high (0.95) for 4000 cells, decreased to 0.61 for 1000 cells, and then to 0.16 for 500 cells. For 25 % doublets, it remained around 0.75 for 1000, 2000, and 4000 cells and decreased to 0.16 for 500 cells.

For S3, 14 SNPs were reported in the complete sample, and the highest decrease from eleven to seven SNPs appeared between 1000 and 500 cells (Fig. 3G). In most downsampled datasets, at least one additional SNP, filtered in the complete sample, was reported, indicating uncertain SNPs with filtering values close to the chosen thresholds. On the complete sample, BnpC inferred three and SCG three/four clusters (8*/*25 % doublets), decreasing to two with fewer cells (<2000) (Fig. 3H). COMPASS, in contrast, inferred eight/nine clusters on the complete sample (8*/*25 % doublets), seven on samples with 1000 and 2000 cells, and three on the samples with 500 cells. With a relatively steady and low number of inferred clusters, SCG’s ARI dropped from an initial 0.80 to 0.60 for 1000 and 0.55 for 500 cells. BnpC inferred the most stable clusters, leading to an ARI above 0.95 for runs with more than 500 cells and an ARI of 0.84 for 500 cells. Relative to the decreasing number of clusters inferred by COMPASS, its ARI decreased from 0.95 to 0.66 for 2000 cells and to 0.53 for 500 cells, for 8 % doublets, and from 0.83 for the complete sample to 0.71 for 2000, 0.62 for 1000, and 0.36 for 500 cells. In contrast to S1 and S2, COMPASS inferred a variable number of clusters on the complete sample (ARI ≠ 1).

In summary, our simulations demonstrated that the number of reported SNPs, inferred clusters, and ARI decreased with fewer cells, as expected. However, the number of inferred clusters remained relatively stable with decreasing sample sizes above 500 cells. BnpC and SCG could distinguish only between a few clusters, if at all, likely clustering the cells into healthy and cancer cells. COMPASS inferred more subtle clonal structures, which remained less well preserved with smaller sample sizes, leading to a higher decrease in its ARI. Additionally, we found no clear trend between the two used doublet rates for SCG and COMPASS.

### 2.4 Successful demultiplexing reveals a high doublet rate

When applied to the multiplexed sample MS1 (Fig. 4B), *demoTape* inferred one cluster with 944 (blue, 22 %), one with 1302 (yellow, 30 %), and one with 966 (green, 22 %) cells. Accordingly, 25 % of the cells were assigned to doublet clusters, marked in Figure 4B by two colors in the bars indicating the cluster assignment. 25 % represents a lower bound for the overall doublet rate, as it does not account for doublets of two cells from the same cluster. By assuming the same doublet rate for cells within clusters, the estimated total doublet rate is around 50 %.

**Figure 4:**
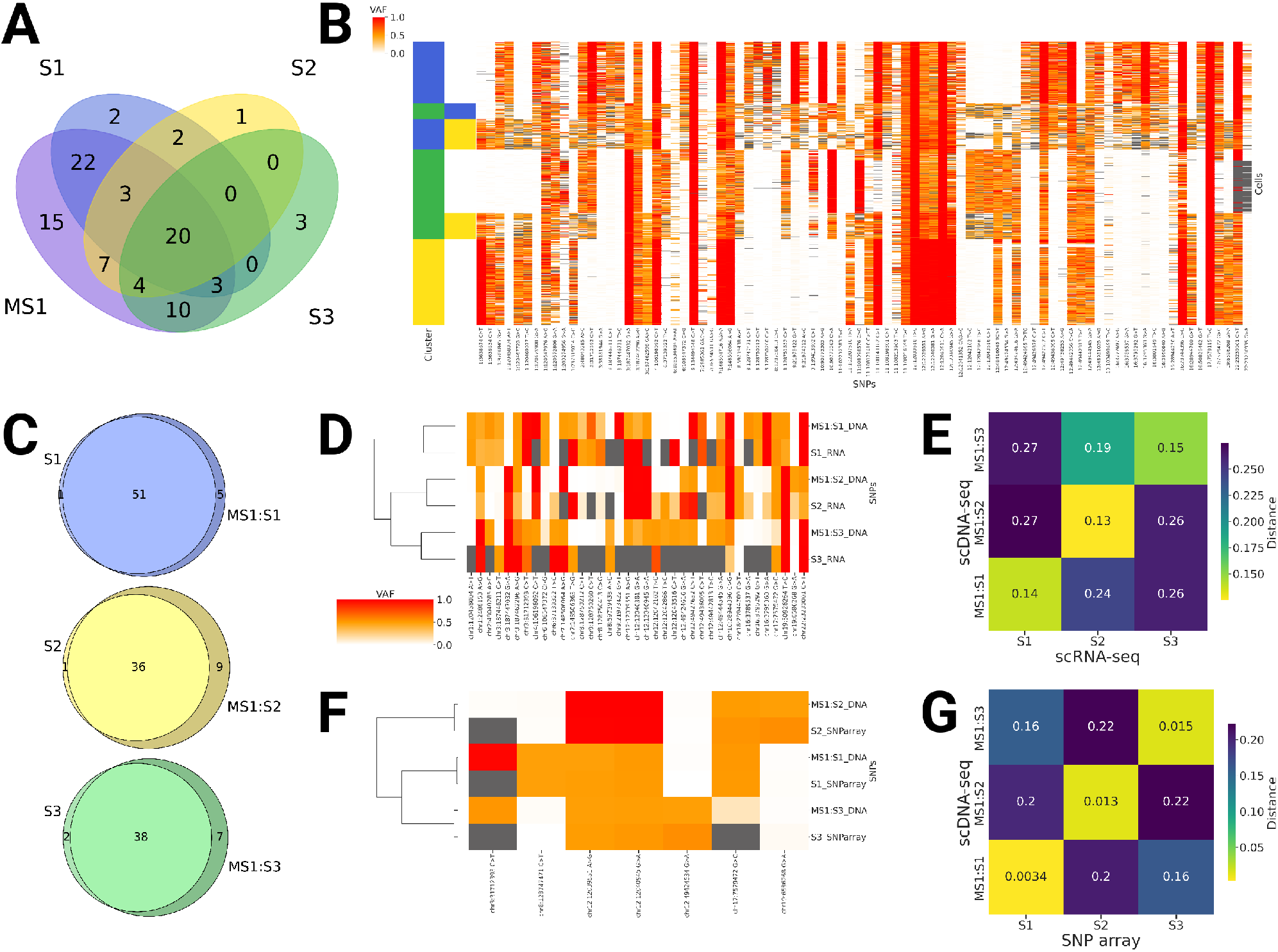
Overview of the multiplexed sample and the demultiplexed cluster-patient assignment. **A** Venn diagram of the SNPs called in the three individually sequenced samples (S1, blue; S2, yellow; S3, green) and the multiplexed sample (MS1, violet). **B** VAF heatmap of MS1. The colored bars on the left indicate the inferred patient and doublet clusters. **C** Venn diagrams of the SNPs called in the three individually sequenced samples and the respective demultiplexed samples MS1:S1, MS1:S2, and MS1:S3. **D** VAF heatmap of the SNPs overlapping between the demultiplexed scDNA-seq clusters (averaged over all cells) and the matched, patient-specific scRNA-seq data (averaged over multiple samples). **E** Pair-wise normalized Euclidean distance between scDNA-seq cluster and scRNA-seq patient SNP profiles per patient. Profiles from the same patient are closer together. **F** VAF heatmap displaying the overlap between the demultiplexed scDNA-seq clusters (average) and the matched, patient-specific SNP array data. **G** Pairwise normalized Euclidean distance between scDNA-seq SNPs and SNP array profiles per patient. Profiles from the same patient are closer together.

Visual inspection of the VAF heatmap showed a clear separation of the data into three blocks, corresponding to the inferred clusters. Especially SNPs called in just one sample correlated clearly with the inferred clusters. For example, at chr1:3638674 (Fig. 4B, first column/SNP), a homozygous SNP (VAF ≈ 1: red) was called in the yellow cluster and a wildtype genotype (VAF ≈ 0: white) in the blue and green clusters. In the doublet clusters containing cells from the yellow clone, heterozygous SNPs (VAF ≈ 0.5: orange) were called in most cells, whereas in the doublet cluster without cells from the yellow clone (i.e., only green and blue cells), wildtype genotypes were called predominantly.

By evaluating the presence of patient-specific SNPs (Fig. 4A; S1: 22 SNPs, S2: 7 SNPs, S3: 10 SNPs) in the inferred clusters in MS1, we identified the blue cluster as S1 (MS1:S1), the yellow one as S2 (MS1:S2), and the green one as S3 (MS1:S3). When comparing the SNPs reported from the individual samples with the demultiplexed ones, most SNPs were detected in both samples (Fig. 4C). Counter-intuitively, more unique SNPs were reported in the demultiplexed samples (5, 9, and 7 for MS1:S1, MS2:S2, and MS3:S3, respectively). The main reason for this is that more SNPs are reported initially in the individual samples, leading to more SNPs within close genomic proximity, which are subsequently filtered as they are likely sequencing artifacts.

### 2.5 SNP profiles enable the assignment of demultiplexed clusters to patients

If multiplexing is applied in practice to save labor and expenses, individually sequenced samples are generally not available. Accordingly, either additional matched SNP data per patient or a combinatorial approach are necessary to assign the demultiplexed clusters back to patients. For matched SNP data, patient-specific SNPs from any source, e.g., from SNP assay, scRNA-seq data, or whole-exome/genome bulk sequencing, can be used as long as enough patient-specific SNPs overlap with the multiplexed data to assign clusters unambiguously to patients. For combinatorial demultiplexing, careful experimental design [29], relying on splitting and multiplexing the samples according to specific patterns, is required. Here, we used patient-specific SNPs derived from (1) SNP arrays and (2) matched scRNA-seq data (see Section 5.6) to assign the demultiplexed clusters to patients.

First, we defined all loci as relevant that were reported after demultiplexing in the individual clusters MS1:S1, MS1:S2, and MS1:S3 or in the multiplexed sample MS1. To obtain the VAF information at all loci, also wildtype ones, we reran the SNP filtering of the demultiplexed clusters but included all relevant loci. Next, we obtained the SNP profiles for the clusters by averaging the VAFs over all cells and excluded SNPs with similar VAF in all scDNA-seq data (VAF *>*95 %, VAF *<*5 %, or VAF 45–55%), as they are uninformative for distinguishing between clusters.

Next, we identified the loci from the SNP arrays and scRNA-seq data that overlap with the cluster SNP profiles, resulting in 37 SNPs overlapping with the scRNA-seq data (Fig. 4D) and seven with the SNP arrays (Fig. 4F). Based on these loci, we calculated the normalized Euclidean distance between between all pairs of SNP profiles and iteratively, starting with the smallest distance, assigned the cluster to the patient SNP profile with the lowest distance, excluding that cluster and patient profile from further assignments afterward.

The SNP profiles derived from scRNA-seq data and SNP arrays were sufficient to assign the demultiplexed samples unambiguously back to patients. The distance matrices of the scRNA-seq data (Fig. 4E) and SNP array (Fig. 4G) profiles showed clearly that cluster MS1:S1 is closest to S1, cluster MS1:S2 closest to S2, and cluster MS1:S3 closest to S3. Even the assignment of cluster MS3:S3 to S3 based on scRNA-seq is distinct, although the scRNA-seq profile of S3 contains 21 missing values.

### 2.6 Comparison of single and multiplexed samples

To compare the called SNPs and downstream analysis between the demultiplexed and the individually sequenced data, we reran the filtering for each demultiplexed cluster separately, filtered uninformative SNPs (Section 5.5), and inferred the evolutionary history of each sample using COMPASS.

The VAF heatmaps of S1 and MS1:1 were similar in large parts: 12 out of 17 informative SNPs reported in S1 were also reported in MS1:S1, and three SNPs detected in MS1:S1 were absent in S1 (Fig. 5A). Accordingly, the inferred event trees also showed similar structures, differing mainly in the placement of SNPs chr6:106534484 and chr15:93545223. In S1, chr6:106534484 was an early cancer event present in all cancer cells, and chr15:93545223 defined a cancer clone (Fig. 5B). In MS1:S1, in contrast, chr6:106534484 was only present in a cancer subpopulation, and chr15:93545223 was present in all cells, hence, likely being a germline event (Fig. 5C). All other differences resulted from SNPs not called in one of the samples. Except for SNP chr6:106534484, both trees contained four cancer clones, and the relative clone sizes were also similar, where applicable (chr16:28944230: 12*/*13 %; chr13:103498499 8*/*8 %; chr6:106534484 20*/*23 % for S1/MS1:S1, respectively).

**Figure 5:**
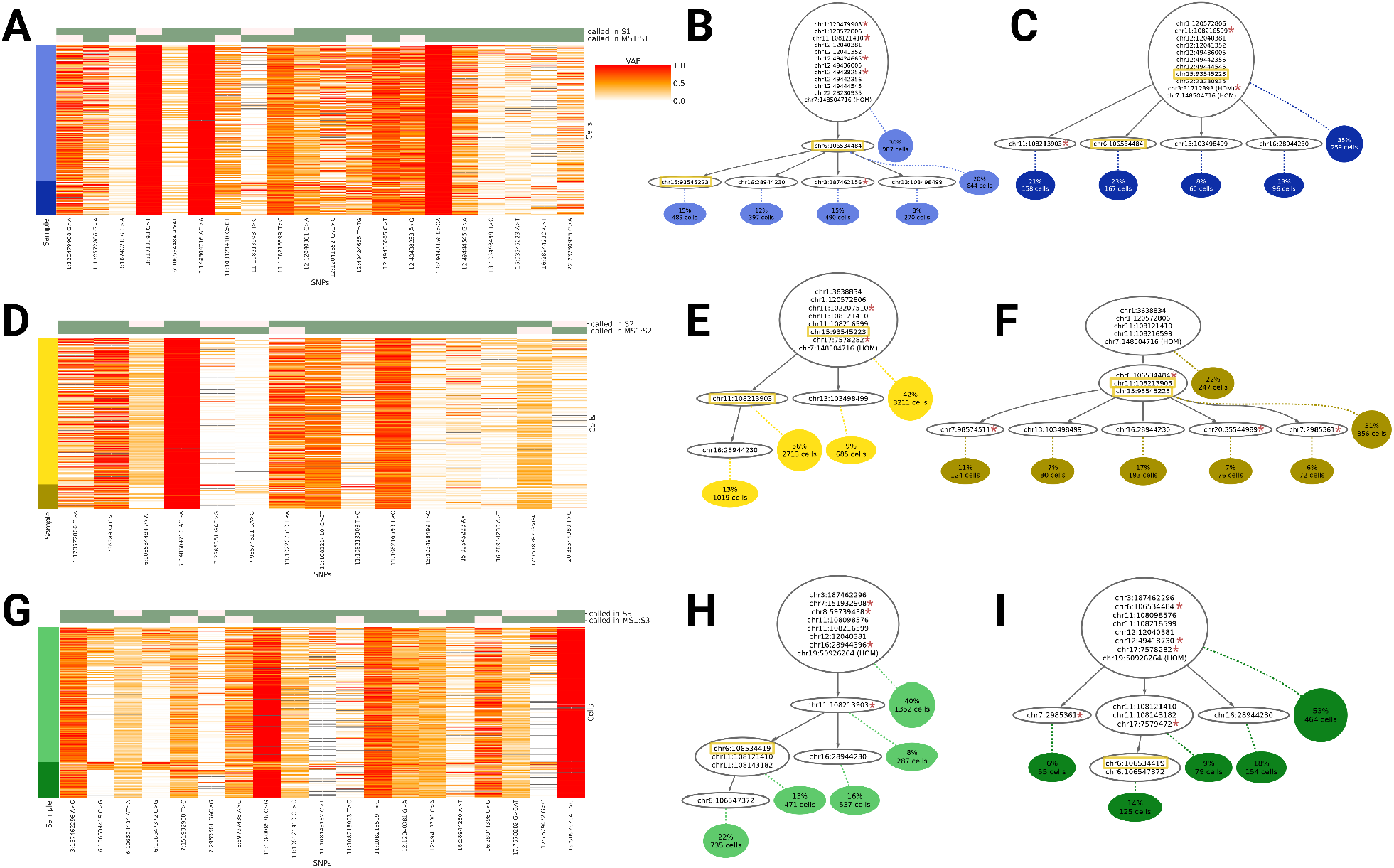
Comparison of the individual samples (S1, S2, S3) with the demultiplexed clusters of the pooled sample (MS1). **A**) VAF heatmap for S1 (light blue) and MS1:S1 (dark blue) and the corresponding event trees for S1 (**B**) and MS1:S1 (**C**). **D**) VAF heatmap for S2 (light yellow) and MS1:S2 (dark yellow) and the corresponding events trees (**E**) and (**F**), respectively. **G**) VAF heatmap for S3 (light green) and MS1:S3 (dark green) and the corresponding event trees **H** and **I**, respectively. The green and red column colors above the VAF heatmaps indicate whether the SNP was reported in the corresponding sample. Event trees were inferred with COMPASS. Yellow boxes indicate SNPs placed at different nodes in the tree topology between the two samples. Red asterisks indicate SNPs reported here but not in the corresponding sample. HOM = homozygous SNP.

In S2 and MS1:S2, we also observed similar genotype heatmaps and tree structures: nine out of eleven informative SNPs found in S1 intersected with the SNPs reported in MS1:2, which contained four informative SNPs not found in S2 (Fig. 5D). The inferred trees showed similar evolutionary trajectories, differing only in the placement of SNPs chr11:108213903 and chr15:93545223, as well as SNPs unique to one of the samples. In S2, chr15:93545223 was inferred as a germline SNP and chr11:108213903 defined a cancer subclone (Fig. 5E), whereas in MS1:S2, both SNPs were initial cancer-forming events present in all cancer cells (Fig. 5F). The size of comparable cancer clones was again similar: 13*/*17 % for chr16:28944230, 36*/*31 % for chr11:108213903, and 9*/*7 % for chr13:103498499, respectively for S2/MS1:S2.

Out of 14 informative SNPs detected in S3, ten overlapped with the SNPs of MS1:S3, and MS1:S3 contained five SNPs not reported in S3 (Fig. 5G). Besides the SNPs unique to only one of the samples, the inferred trees varied only in the SNP chr6:106534419. In S3, it was placed in the same event node with two SNPs on chromosome 11 (Fig. 5H), whereas in MS1:S3, the two chromosome-11 SNPs occurred before chr6:106534419 (Fig. 5I). The only node directly comparable in size contained the SNP chr16:28944230, where the relative sizes were 16*/*18 % for S3/MS1:S3.

Overall, the SNPs and trees of the individual and demultiplexed samples overlapped widely for all three samples. The individually sequenced and multiplexed samples missed a similar amount of informative SNPs present in their counterpart, the inferred trees shared a similar structure, and comparable clones were alike in size. It is apparent from the VAF heatmaps that differences in the reported SNPs originated not from technical variations but from the filtering steps, as all SNP loci were covered in the individually sequenced and the demultiplexed samples. Instead, the differences resulted from SNPs classified as uninformative germline SNPs in one of the samples but passed the filtering thresholds in the other. Visual inspection of the VAF heatmaps and the inferred trees revealed that not all germline SNPs were filtered (e.g., Fig. 5A: chr7:148504716).

## 3 Discussion

Recent technological advances enable the DNA sequencing of thousands of individual cells, providing a snapshot of the DNA of the sequenced tissue at single-cell resolution. However, scDNA-seq is still labor- and cost-intensive, hampering its widespread use. In this work, we introduce *demoTape*, a computational distance-based demultiplexing approach for targeted scDNA-seq data. Multiplexing allows the preparation and sequencing of multiple samples jointly, decreasing the expenses and working hours required per sample. First, we sequenced three samples individually and used these to simulate datasets with known ground truth. *DemoTape* outperformed state-of-the-art scRNA-seq methods and demultiplexed the cells with high accuracy. When downsampling the number of cells in each of the three individual samples, the clustering consistency decreased for the three applied clustering algorithms, as expected. However, the number of SNPs, clusters, and ARI remained relatively constant for samples with at least 1000 cells and decreased highest from 1000 to 500 cells. Accordingly, these findings suggest avoiding multiplexing samples such that the number of cells per sample drops to 500. With the currently achieved average cell numbers of 5000 to 6000 cells per run [30, 31] and doublet rates between 8 and 25 %, this suggests that three or four samples can readily be multiplexed, but multiplexing more than five samples may not be advisable.

We multiplexed, sequenced, and analyzed the three samples jointly. Surprisingly, the demultiplexing revealed that 25 % of the non-empty sequenced droplets were doublets with cells from different samples. By assuming the same rate for doublets within each sample, the total doublet rate would be around 50 %. Next, we successfully assigned the inferred clusters to the original samples, using SNPs called in matched scRNA-seq samples or SNP-array data, demonstrating the practicality of the multiplexing approach. Finally, we compared the clonal compositions and inferred SNP trees of the demultiplexed samples with the individual ones, finding a high consistency between the genotypes and tree structures for all three samples. However, more precise filtering of germline SNPs while retaining somatic SNPs remains an open challenge, as VAF distributions were rather noisy, with varying noise levels per SNP/amplicon.

Similar to approaches for the computational demultiplexing of scRNA-seq data like souporcell or scSplit, *demoTape* is based on the data-inherent SNPs and renders additional biochemical assays redundant. However, the underlying input data varies greatly: scRNA-seq data does not amplify the sequenced molecules and covers only SNPs in expressed genes, making it sparse and noisy. Additionally, the widely used 10x Genomics protocols sequence only the 3’ or 5’ ends of genes, leading to even sparser data. scDNA-seq data, in contrast, is smaller in size and covers most target sites, resulting in distinct genotype profiles per cell and sample. Using our proposed cell distance, these can be used to calculate distances between SNP profiles, enabling accurate and fast demultiplexing of targeted scDNA-seq data. Cardielloe *et al*. [32] recently showed that genotype-based demultiplexing of scRNA-seq data works for non-human organisms, and so should also apply to scDNA-seq with even higher accuracy. The expansion of genotype-based demultiplexing of single cells to species beyond humans and data modalities beyond RNA enables the investigation of broader questions requiring large amounts of data.

The high number of doublets led to a loss of 25 % of the data, which, depending on the number of sequenced cells and the mixing ratio, could bias the downstream analysis and hamper accurate inference of the clonal composition. Given that we know the clonal composition of the individual samples (e.g., through clustering methods like the ones applied here), doublets could be split and reassigned to clones from different samples. However, this would correct the size of the inferred clones only; clones missed due to small size could not be recovered by this approach. When run multiple times on the complete sample, SCG and BnpC inferred a low number of clusters, likely distinguishing only between healthy and cancer cells, and these clusters were also inconsistent for two out of three samples. COMPASS inferred more clusters, possibly identifying cancer subpopulations better, but in one out of three samples, the inferred clusters were also inconsistent between runs on the complete sample. These findings suggest that methods designed especially for targeted scDNA-seq data are superior to general ones but still leave room for improvement, and further method development remains highly desirable. For example, one could use the noise profiles from our multiplexing experiment, where it is easy to distinguish between true SNPs and background noise based on the individually sequenced samples, and integrate these rates into the clustering methods. It became apparent that the filtering thresholds in the data processing play a crucial role in the quality and quantity of the reported SNPs. For downsampled and demultiplexed samples, for example, to account for the smaller sample size, we changed the minimum frequency for SNP loci to be reported (from 1% of cells to 50 cells). A systematic evaluation of the filtering thresholds remains open. Our study suggests that the data quality would benefit from fine-tuned thresholds and additional filters, likely also easing downstream analysis and preventing false positives, easing the future use of targeted scDNA-seq.

## 4 Conclusions

In summary, we introduced *demoTape*, an accurate demultiplexing approach for targeted scDNA-seq data based on the distance between cellular VAFs. When applied, *demoTape* revealed that the clonal composition is mainly retained despite the lower cell number. Multiplexing without additional biochemical assays increases throughput and reduces the working time and costs required per sample, possibly promoting its broader use. Especially for genetic diseases like cancer, more DNA-seq data at single-cell resolution offers great potential to understand the mechanisms underlying carcinogenesis and to adjust patient treatment according to personal genotype profiles.

## 5 Methods

This section describes the demultiplexing algorithm and the simulation experiments in detail, followed by an overview of the patient samples obtained and their processing and a summary of the software implementation.

### 5.1 Computational demultiplexing of samples

Given a sequenced and processed multiplexed sample containing cells from *s* individual samples, for demultiplexing *demoTape* requires the read count matrix 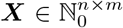, where *n* is the number of cells and *m* is the number of SNP sites. Each *x*_*ij*_ = (*a*_*ij*_, *r*_*ij*_, *t*_*ij*_) contains the read count *a*_*ij*_ supporting the alternative allele, the read count *r*_*ij*_ supporting the reference allele, and the total read count *t*_*ij*_ = *a*_*ij*_ + *r*_*ij*_ in cell *i* at site *j*. ***X*** can be generated from the Loom or H5 files that are part of the Tapestri pipeline output after filtering these with the Mosaic software package [33].

By modeling the read counts with a binomial distribution and assuming that they are i.i.d. per site, we can define the likelihood for cell *i* as

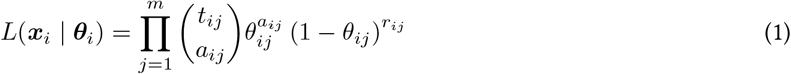

The likelihood is maximal at

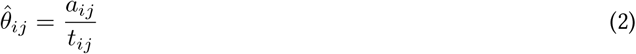

where 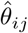 is the observed variant allele frequency (VAF) in cell *i* at site *j*.

As distance between two cells *i* and *i*^*′*^ we define their log-likelihood ratio in a separate versus a joint binomial model. Specifically, we consider the ratio between the maximum likelihood given that the read counts per cell are samples from independent VAFs 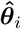 and 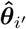, and the maximum likelihood give that the read counts for both cells are sampled from a joint VAF 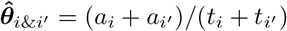. This quantity is small if 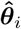 and 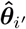 are similar and increases with differences in the VAFs. The log-likelihood distance is defined as

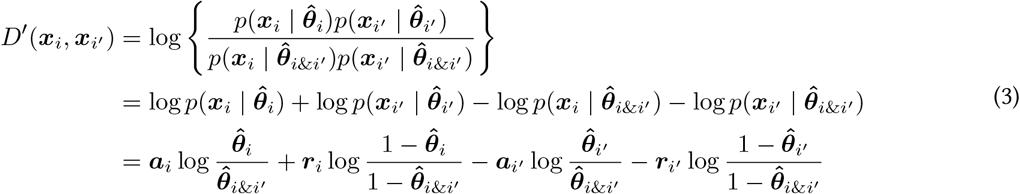

To normalize the distance to values between 0 and 1, we divide it by the maximal possible distance between two cells

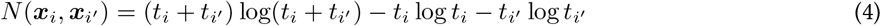

where all reads in one cell support the alternative while all reads in the other cells support the reference allele, or vice versa. Accordingly, the normalized distance between two cells is

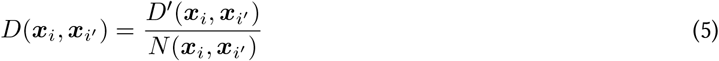

*DemoTape* calculates the pairwise distance between all cells and applies hierarchical agglomerative clustering (HAC) with Ward’s linkage [34] to generate a dendrogram. Next, *demoTape* cuts the dendrogram such that 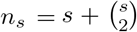 clusters ***k*** = {1, 2, …, *n*_*s*_} are defined, representing *s* individual samples and 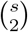 doublet clusters. The doublet clusters contain sequences obtained from two distinct samples, i.e., two cells encapsulated in the same droplet due to a technical error. The generated cluster assignment can be represented as a vector ***c*** ∈ ***k***^*n*^, where *c*_*i*_ is the assigned cluster of cell *i*. To identify which clusters correspond to doublets and which to individual samples, *demoTape* defines a cluster profile matrix 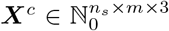. First, the average read depth and VAF weighted by the read depth is calculated for each cluster *k* at each position *j*:

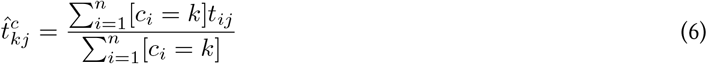

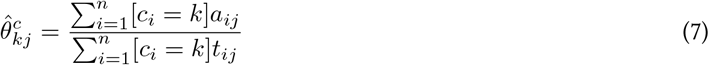

where [·] is the indicator function, which is 1 if the expression inside the brackets is true and 0 otherwise. Using these values, the cluster profiles are then defined as

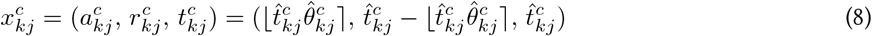

where ⌊·⌋ is the result of rounding to the nearest integer.

Next, *demoTape* creates profiles of all 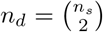 hypothetical doublet clusters ***d*** = (1, 2), (1, 3), …, (*n*_*s*_ − 1, *n*_*s*_)}, where 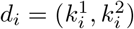 indicates the two clusters 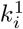 and 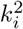 forming the hypothetical doublet cluster *d*_*i*_. Doublet cluster profiles 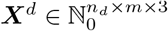 are computed according to equations 6, 7, and 8 by averaging over all cells assigned to the two clusters *d*_*i*,1_ and *d*_*i*,2_.

Given the *n*_*s*_ inferred cluster profiles and the *n*_*d*_ hypothetical doublet cluster profiles, *demoTape* calculates the distance matrix 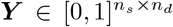 between the profiles using Equation (5), where *y*_*i,j*_ is the distance between the cluster profile 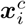 and the hypothetical doublet cluster profile 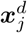. Distances between cluster *i* and hypothetical doublet clusters including *i*, e.g., between cluster 1 and the hypothetical doublet cluster (1, 2), are set to 1 as doublet clusters cannot include themselves. Additionally, ***y***· _,*j*_ is set to 1 if the distance between clusters *d*_*j*,1_ and *d*_*j*,2_ is very small (<0.05), indicating that *d*_*j*,1_ and *d*_*j*,2_ are clones from the same sample, inferred through an insufficient cutting of the dendrogram. Accordingly, *demoTape* can only demultiplex samples that differ by an average distance greater than 0.05, a hyperparameter changeable by the user, as its unable to differentiate between samples and clonal structures within a sample otherwise.

*DemoTape* then identifies *n*_*d*_ doublet clusters according to the following procedure (Algorithm 1): First, *demoTape* takes the smallest distance 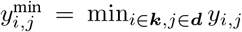 between the cluster and hypothetical doublet cluster profiles, indicating that the inferred cluster *k*_*i*_ is likely a doublet cluster composed of clusters *d*_*j*,1_ and *d*_*j*,2_. To assess if this is true or if the tree cutting generated insufficient clusters, e.g., due to a high distance between two clones within a sample, *demoTape* investigates the homozygous SNP and wildtype loci in the three considered clusters, as these are particularly informative about true doublets (see Section 2.4). Given a VAF cutoff *α* to define homozygous and wildtype loci, e.g., *α* = 0.05, *demoTape* defines loci in individual sample clusters as homozygous if the VAF is ≥1 −*α* and as wildtype if the VAF is ≤*α*. Accordingly, loci in doublet clusters are defined as homozygous if the VAF is ≥1 −*α*^2^ and as wildtype if the VAF ≤ *α*^2^. Wildtype and homozygous SNP loci in a doublet cluster require also wildtype/homozygous SNP loci in both individual sample clusters, respectively, since reads from both samples are aggregated and would otherwise lead to an intermediate VAF. Similarly, homozygous SNPs in just one individual sample cannot result in a homozygous SNP in the doublet cluster. Therefore, *demoTape* counts if all wildtype/homozygous SNP loci in the doublet cluster are also wildtype/homozygous SNP loci in the two individual sample clusters, if all homozygous SNP loci in both individual sample clusters are also homozygous in the doublet cluster, and if loci that are homozygous in just one individual sample cluster are neither a wildtype nor a homozygous SNP in the doublet cluster. If this is true for all but maximum one site, allowing some uncertainty, *demoTape* assumes *k*_*i*_ to be a true doublet cluster (function HomozygousSitesMismatch in Algorithm 1). In that case, *demoTape* removes the rows *i, d*_*j*,1_, and *d*_*j*,2_, and all columns where *d·* _,1_ = *k*_*i*_ or *d ·*_,2_ = *k*_*i*_ from ***Y***, and proceeds identifying the next doublet. In case of insufficient clusters, *demoTape* increases the number of clusters generated through dendrogram cutting by one, recalculates ***Y*** accordingly, and repeats the procedure.

If the number of clusters was not increased, the remaining *s* clusters are the demultiplexed samples. Otherwise, an additional step is required to merge single clusters until *demoTape* obtains *s* clusters. For this, *demoTape* merges the two clusters with the closest distance whose merging does not conflict with the inferred doublet clusters. A conflict exists if any *d*_*i*,1_ = *d*_*i*,2_ or *d*_*i*_ = *d*_*j*_ through the merge, i.e., if a doublet becomes a single cluster or two doublet clusters become identical. If *demoTape* cannot merge two clusters without a conflict, it merges the two closest single clusters and removes the obsolete doublet cluster.

### 5.2 Simulation of multiplexed samples

To generate multiplexed samples with known ground truth, we sampled cells without replacement from the patient samples S1, S2, and S3 according to the mixing ratio. If sites were not sequenced in one or two patients, we set their genotypes to missing values (3) in the corresponding cells. In all simulation runs, we sampled (1 + *δ*) × 3677 cells, where 3677 is the smallest number of sequenced cells in S1, S2, and S3, and *δ* is the doublet rate to simulate. To generate doublets, we chose two patients with probabilities proportional to the mixing ratio and subsampled one cell per patient from the previously sampled cells. For each doublet, we generated the data per locus as following: If the same genotype was called in both cells, we assigned this genotype to the doublet. If one cell had a missing value, we assigned the genotype of the other cell. In any other case, we assigned a heterozygous genotype. As total read depth, alternative allele read depth, reference allele read depth, and genotype quality (GQ), we took the rounded mean of the two cells. Subsequently, we applied the Mosaic filters on the subsampled cells and doublets, treating them as one sample.

#### Algorithm 1 Identification of doublet clusters

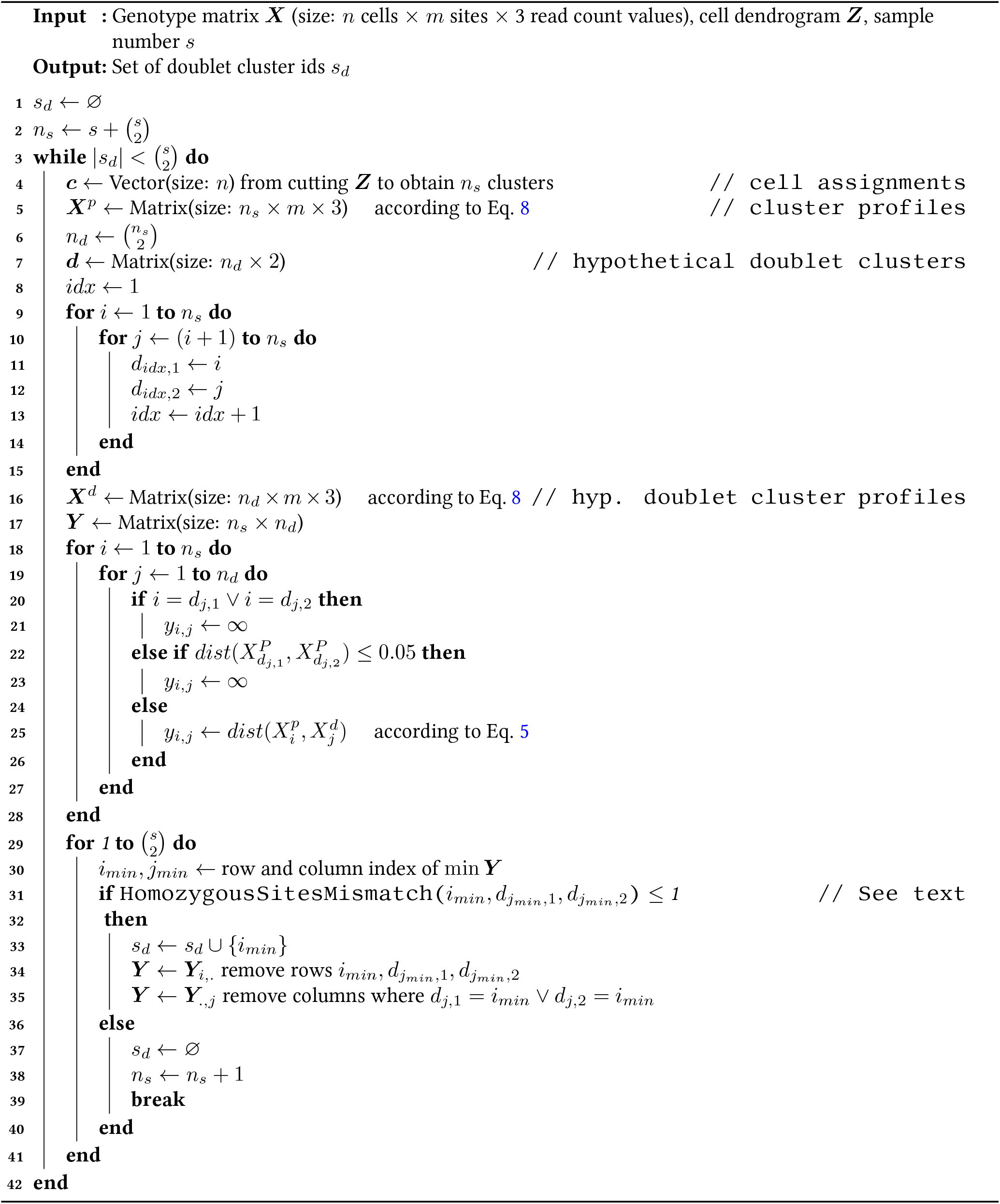

### 5.3 Patient sample collection

Patients with diffuse large B-cell lymphoma were recruited at the University Hospital Zurich and voluntarily provided informed consent to participate in the study without receiving compensation (Ethical Approval: Zurich, Switzerland, 2019-01744). The samples used in this study consisted of sections obtained from fresh-frozen lymph node biopsies from the diagnostic procedure.

### 5.4 Targeted single-cell DNA sequencing

Cells of three samples were thawed, resuspended and washed with RPMI (Gibco) supplemented with 10 % fetal bovine serum (Gibco). Debris and aggregates were removed using a 40 µ*m* cell strainer (Corning) and cells were counted with a K2 Cellometer and ViaStain AOPI Staining solution (Nexcelom Bioscience). 0.5 to 1 ×10^6^ cells per sample were pooled, pelleted and resuspended in 1 *mL* CSB (Cell Staining Buffer, Biolegend). After an additional centrifugation step, cells were resuspended in 50 µ*L* CSB and counted (Nexcelom Bioscience K2 Cellometer). 1 to 1.5 × 10^6^ cells were then labeled with TotalSeq-D Heme Oncology Cocktail (Biolegend) according to the Mission Bio Tapestri protocol with the following adjustments: The lyophylized TotalSeq-D antibodies were resuspended in 120 µ*L* CSB. 50 µ*L* of the reconstituted TotalSeq-D cocktail were added to the blocked cell pool. After labeling, the cells were washed three times in 14 *mL* and two times in 1 *mL*, and finally filtered (40 µ*m* cell strainer, Corning).100 × 10^3^ cells were used for encapsulation in the Tapestri instrument and further processed for single-cell DNA-seq according to Mission Bio’s guidelines (v2 reagents). Tapestri single-cell DNA libraries were sequenced with an Illumina NovaSeq 6000 system. The target panel of 533 amplicons, covering 99 genes, was designed to capture known SNPs and copy-number-variations across a wide set of B-cell lymphoma types and implemented with Mission Bio’s Tapestri Designer.

### 5.5 Single cell DNA data processing

We processed all samples with the Mission Bio Tapestri DNA pipeline (v2.0.2) with default parameters and filtered the data using Mosaic with default parameters. After demultiplexing MS1, we selected the cells assigned to each cluster and applied the Mosaic filtering on only these cells with default parameters, except for the minMutated filter, which we set to an absolute value of 50 cells instead of 1 % of the cells. We also set the minMutated filter to 50 cells for the downsampling simulations, due to the decreased sample size.

For the comparison between the multiplexed and individually sequenced samples and for the downsampling simulations, we filtered the inferred SNPs further to obtain only SNPs informative of the cancer, i.e., exclude germline SNPs and technical artifacts. If in 99% of all cells the VAF was >0.95 or <0.05, SNPs were considered as homozygous germlines or false positives, respectively. To exclude heterozygous germline SNPs or technical artifact, we assume that their VAF is symmetrically distributed around 0.5. We tested for symmetry with a FWER corrected two-sided Kolmogorov–Smirnov test (*α* = 0.01), taking all VAF values 0 *<* VAF *<* 0.5 as the first distribution, and 1 −(0.5 *>* VAF *>* 1) as the second distribution.

For the clustering analysis, we ran COMPASS v1.1 (--nchains 1 --chainlength 50000 –CNV 0 --doubletrate *δ*), BnpC v1.0 (-n 1 -r 120 -pp 1 1 -ap 1 1 -cup 0), and SCG (py3 branch: num_clusters: 20, model: doublet, alpha_prior: [(1-*δ*)/*δ*, 1]; --max-iters 100000). For the comparison between the multiplexed and individually sequenced samples, we ran COMPASS with parameters --nchains 4 --chainlength 50000 --CNV 1 --doubletrate 0.25. To visualize trees inferred with COMPASS, we concatenated low-frequency nodes with less than 5 % of cells attached by excluding low-frequency leaf nodes and concatenating internal nodes with just one outgoing edge with their descending nodes. If neither was applicable for a low-frequency node, we concatenated it with its parent node if the parent had just one outgoing edge (i.e., the low-frequency node was the only child node) or retained the node otherwise.

### 5.6 Single-cell RNA sequencing and data processing

Cells were thawed, resuspended and washed with RPMI (Invitrogen) supplemented with penicillin/streptomycin (Invitrogen), L-glutamine (Invitrogen), and 10 % fetal calf serum (FCS, Gibco) (RPMI-FCS) to remove DMSO. Fc-receptors were blocked by adding Human TruStain FcX (Biolegend). Cells were then stained with TotalSeq-C (Biolegend) hash antibodies (1.25 µ*g mL*^*−*1^) for 30 *min* at 4 ^*°*^*C*, washed 3x in Cell Staining Buffer (Biolegend). Cells were then pooled and stained with TotalSeq-C Human Universal Cocktail (Biolegend) for 30 *min* at 4 ^*°*^*C*, washed 3x in Cell Staining Buffer and processed using the 5’ HT Single Cell GEX and VDJ v2 platform (10x Genomics) according to manufacturer’s recommendations. Sequencing libraries were prepared following the manufacturer’s recommendations (10x Genomics). Final libraries were pooled and sequenced on an Illumina NovaSeq NovaSeq 6000 system, followed by demultiplexing, FASTQ conversion, and alignment to GRCh38 with the Cell Ranger multi pipeline (v7.0.0, 10x Genomics). Cells with fewer than 500 UMIs, fewer than 250 genes, more than 10 % mitochondrial UMIs, with failed antibody library preparation (0 antibody UMI counts in more than 50 % of antibodies), or cells that were part of antibody aggregates (isotype control counts above 3 median absolute deviations) were excluded. Doublets were removed based on three complementary approaches; hash-oligos (hashedDrops() from scran [35]), SNP profiles and doublet simulation (scDblFinder [36]). Finally, we clustered cells and called SNPs per cluster using souporcell [22] (v2.0), and used Picard’s [37] LiftoverVcf to lift the genome coordinates of the RNA SNPs to GRCh37, matching the DNA coordinates. The scRNA-seq patient data was obtained from multiple sequencing runs with different patients multiplexed in each: S1 was included in one, S2 in eleven, and S3 in two multiplexed sequencing runs.

### 5.7 Implementation

*DemoTape* is implemented in Python and requires the read counts at all target loci in Loom file format and the number of multiplexed samples as input. Additional SNP profiles obtained from scRNA-seq, SNP assays, or WES/WGS DNAseq are required to assign the demultiplexed clusters to patients. The pipelines for simulating and downsampling data, and a full analysis and demultiplexing pipeline that takes as input the loom files generated by the Mission Bio Tapestri DNA pipeline, were implemented in Snakemake.

## 6 Data and code availability

scDNA-seq and scRNA-seq data of the three lymphoma patients are available upon request from according to the data sharing policy of the INTeRCePT consortium (in preparation by the consortium).

All original code has been deposited at https://github.com/cbg-ethz/demoTape and is publicly available.

## Author contributions

Conceptualization, N.Be., J.K, and A.M.; Methodology, N.Bo.; Software, N.Bo.; Validation, N.Bo. and J.G.; Formal Analysis, N.Bo.; Investigation, C.B., T.D., M.J.F, M.P., M.R., and S.R.; Resources, C.B., N.Be., M.R., and T.Z.; Data Curation, N.Bo. and J.G.; Writing - Original Draft, N.Bo.; Writing - Review & Editing, N.Be., N.Bo., J.G., J.K. and M.P.; Visualization, N.B.; Supervision, N.Be.; Funding Acquisition, N.Be. and T.Z.; Project administration: T.Z.

## Acknowledgments

This work was supported by the European Union’s Horizon 2020 Research and Innovation Program under the Marie Sklodowska-Curie CONTRA grant agreement No. 766030, the European Research Council (ERC) agreement No. 609883, and the LOOP Zurich INTeRCePT project.

